# DPAM: A Domain Parser for AlphaFold Models

**DOI:** 10.1101/2022.09.22.509116

**Authors:** Jing Zhang, R. Dustin Schaeffer, Jesse Durham, Qian Cong, Nick V. Grishin

**Author notes:** These authors contributed equally (Jing Zhang, R. Dustin Schaeffer, Qian Cong, Nick V. Grishin).

## Abstract

The recent breakthroughs in structure prediction, where methods such as AlphaFold demonstrated near atomic accuracy, herald a paradigm shift in structure biology. The 200 million high-accuracy models released in the AlphaFold Database are expected to guide protein science in the coming decades. Partitioning these AlphaFold models into domains and subsequently assigning them to our evolutionary hierarchy provides an efficient way to gain functional insights of proteins. However, classifying such a large number of predicted structures challenges the infrastructure of current structure classifications, including our Evolutionary Classification of protein Domains (ECOD). Better computational tools are urgently needed to automatically parse and classify domains from AlphaFold models. Here we present a Domain Parser for AlphaFold Models (DPAM) that can automatically recognize globular domains from these models based on predicted aligned errors, inter-residue distances in 3D structures, and ECOD domains found by sequence (HHsuite) and structural (DALI) similarity searches. Based on a benchmark of 18,759 AlphaFold models, we demonstrated that DPAM could recognize 99.5% domains and assign correct boundaries for 85.2% of them, significantly outperforming structure-based domain parsers and homology-based domain assignment using ECOD domains found by HHsuite or DALI. Application of DPAM to the massive set of AlphaFold models will allow for more efficient classification of domains, providing evolutionary contexts and facilitating functional studies.

## Introduction

Annotation of proteins with their constituent domains is a fundamental step towards understanding their evolution and function. Protein domains are conserved regions conveying evolutionary fitness through function[1]. Partitioning a protein sequence into domains and classifying each domain into evolutionarily related groups is an essential step in functional annotation. The predicted function can be used to interpret high-volume data from large-scale studies. For poorly understood proteins, homology to other domains helps to generate experimentally testable hypotheses, and accurate domain boundaries are essential for designing gene constructs for experimental studies[1-4]. To date, resources for the classification of protein domains have fallen into two groups: sequence classifications that classify according to amino acid sequence, such as Pfam and CDD[5, 6] and structure classifications that are largely based on experimentally determined spatial structures, such as SCOP[7] and CATH[8]. Structural classifications identify remote homologs and possess accurate domain boundaries but have been constrained to the small fraction of proteins with experimental 3D structures.

Our Evolutionary Classification Of protein Domains (ECOD), is a hierarchical domain classification of protein structures specifically tailored to identifying and classifying distant evolutionary relationships, i.e., remote homology[9]. Domains sharing homology deduced from sequence and profile searches or revealing structural similarity coupled with functional evidence are grouped into ECOD H-groups. The current ECOD release (v285) consists of over 890,000 domains in 3,715 H-groups derived from nearly 600,000 peptide chains from over 180,000 structural depositions in PDB[10, 11]. ECOD has been accepted as a standard by the field: (1) every PDB entry is linked to the ECOD classification to provide the evolutionary context; (2) ECOD is incorporated into software tools, such as HHsuite[12] and RUPEE[13], as a search database; (3) ECOD serves as the source of homologous domains for target classification in multiple rounds of Critical Assessment of techniques in protein Structure Prediction (CASP), a community-wide experiment for structure predictors to test their methods against target sequences whose structures are not yet public.

The latest round of CASP[14, 15] (CASP14) revealed a breakthrough in the structure prediction field: AlphaFold (AF), developed by DeepMind, predicted 3D structures of proteins from their sequences with accuracies approaching those of experimental structures, including numerous difficult examples of fast-evolving proteins[6, 16]. DeepMind has been using AF to model proteins of biomedical importance, and in partnership with European Molecular Biology Laboratory, they released 3D structures for over 200 million proteins from model organisms, human pathogens, and representative UniProt entries in the AlphaFold protein structure DataBase (AFDB)[17]. This breakthrough is transforming structural biology, where computation is becoming a key component in solving 3D structures of the most challenging and important protein complexes[18, 19] and designing small molecule drugs to target specific structures[20, 21].

This breakthrough in structure prediction is expected to guide the course of protein science in the near future by speeding up the discovery and characterization of proteins with novel and important functions. To maximize the benefits of these predicted structures, it is essential to partition them into domains, evaluate their quality, and classify them by their evolutionary relationships. However, the massive number of AlphaFold models represents a challenge for structure classifications, including ECOD, which are currently only adapted to classify experimental structures. ECOD is frequently updated to include newly released experimental structures. These updates are done through a combination of automatic assignment with human expert curation. The current ECOD automatic domain assignment pipeline starts from BLAST searches against ECOD domains and previously classified PDB chains to identify homologs with high sequence similarity^[22]^. Subsequently, distant sequence hits identified by HHsuite against a set of domain profiles are used to assign regions that cannot be assigned by BLAST. Then, the structural domain parser PDP is used to make small alterations to domain boundaries^46^. Finally, after automatic determination of non-domain regions, unassigned domains (5% − 10% cases) are subject to manual curation.

Incorporating the large number of AlphaFold models into structure classifications faces several challenges. First, AF models contain a significant fraction of regions not suitable for globular domain classification, including disordered segments (**Figure 1A**), single transmembrane helices, signal and other protein sorting peptides, linkers between globular domains, and coiled coils (**Figure 1B**). We refer to these regions as ‘non-domain regions’ for simplicity. Structure similarity in such regions frequently arises from convergence instead of homology. Therefore, annotating the non-domain regions is an essential task. Second, structure classifications databases principally rely on manual curation to confirm structure similarity, which cannot be readily scaled up to hundreds of millions of AlphaFold models. Thus, better computational tools to recognize globular domains and to facilitate automatic domain classification are necessary.

**Figure 1.**
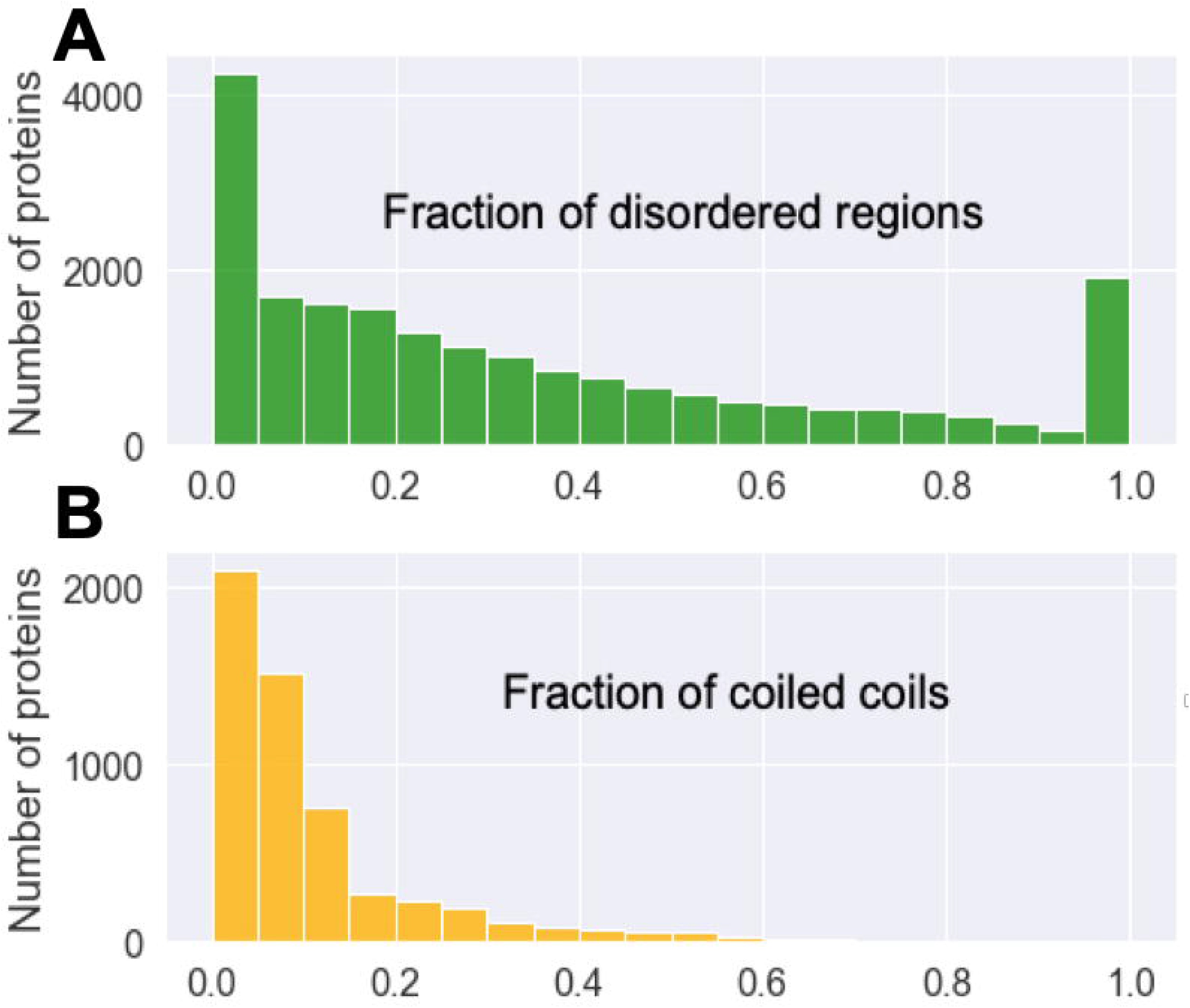
**The fraction of (A) disordered regions** (predicated by predicted aligned errors of AlphaFold models) **and (B) coiled-coils** (predicted by NCOILS[31]) **in AlphaFold models of human proteins**. Proteins without coiled-coils are not included in **B**.

Here, we present a domain parser to recognize single globular domains from AlphaFold protein models. Our Domain Parser for AlphaFold Models (DPAM) combines several types of evidence, including the Predicted Aligned Errors (PAE) associated with each AlphaFold model, the residue-residue distances in AlphaFold model, and homology-based domains using candidate homologous ECOD domains detected by HHsuite [23] and DALI [24]. Our benchmark based on previously classified proteins in ECOD shows that this domain parser can recognize 99.5% of these domains, and the boundaries of 85.2% of these domains agree with the ECOD definitions. Such performance is around two times better than the previous structure domain parsers, PDP [25] and PUU [26]. Once an AlphaFold model is partitioned into single domains, assigning the domains to evolutionary hierarchy becomes much straightforward. Therefore, we expect this tool to be broadly useful for researchers who are interested in analyzing AlphaFold models.

### Develop a benchmark set for parsing domains in AlphaFold models

To benchmark our domain parser, we needed a set of correctly parsed domains, and we derived such a benchmark set from the overlap between proteins that were included in AFDB and those whose experimental structures were already classified in ECOD. Until June 2022, AlphaFold Database contained 992,000 models that covered proteins from model organisms and human pathogens as well as reviewed entries from Uniprot. We obtained the 3D structure and PAE plots for each of these proteins from AFDB. We extracted the sequences of all ECOD domains (v285) (http://prodata.swmed.edu/ecod/complete/distribution). We searched for ECOD domains homologous to each AlphaFold model by DIAMOND [27]. 585,000 AlphaFold models show close homology (e-value < 0.0001) to ECOD domains, and 87,000 were at least partially classified by ECOD (i.e., >= 95% sequence identity to ECOD domains) because 3D structures of some domains in these proteins were solved in experimental structures.

After removing redundancy by mmseqs2 [28] (identity >= 50%, coverage > 80%) and mapping these 87,000 proteins to ECOD, we obtained 18,759 AlphaFold models that were used as the benchmark set. 10,545 (56%) proteins from the benchmark were completely classified in ECOD, and the rest were partially classified because the experimental structure did not cover the entire protein. 6,776 (36%) proteins in this benchmark contain multiple classified ECOD domains, and an additional 4,714 (25%) proteins likely contain multiple domains based on our domain parser (detailed **below**), but the current ECOD only classified a single domain for each of them. This dataset was used to test the performance of our method, and it is available at https://github.com/CongLabCode/DPAM.

### Prepare data for parsing domains

We obtained the PDB70 (non-redundant PDB entries filtered by 70% sequence identity) database for HHsuite from http://wwwuser.gwdg.de/~compbiol/data/hhsuite/databases/hhsuite_dbs/. We parsed the PDB70 database to obtain each of the 92,111 representative PDB chains and further found 126,416 ECOD domains defined in these chains. We further removed redundancy in these ECOD domains by mmseqs2 (identity >= 70%, coverage > 80%), and a total of 63,065 representative ECOD domains were selected as a result. These ECOD domains were included in a database for DALI search, which we dubbed the ECOD70 database.

For each AlphaFold model in the benchmark set, we identified its sequence hits in the PDB70 database by HHsuite. Since the vast majority of PDB entries were classified in ECOD, based on these classifications, we partitioned each PDB70 hit into domains, and thus the AlphaFold models were indirectly mapped to ECOD domains. In addition, we identified the structural hits for each AlphaFold model by DALI search against the ECOD70 database. Although DALI remains the best tool to find structural similarities [24], in most cases, it aligns an ECOD domain to only one segment in a query structure even if the query contains multiple copies of this ECOD domain. Thus, DALI cannot detect duplicated domains in proteins. To alleviate this problem, we developed an iterative DALI alignment procedure (**Figure 2E**) for ECOD hits found by a traditional DALI search run. In iterations, the segment of a query protein that was aligned to an ECOD domain in a previous round, is excluded in the next round of DALI search, until no similarity is found between the remaining portion of the query and the ECOD domain.

**Figure 2.**
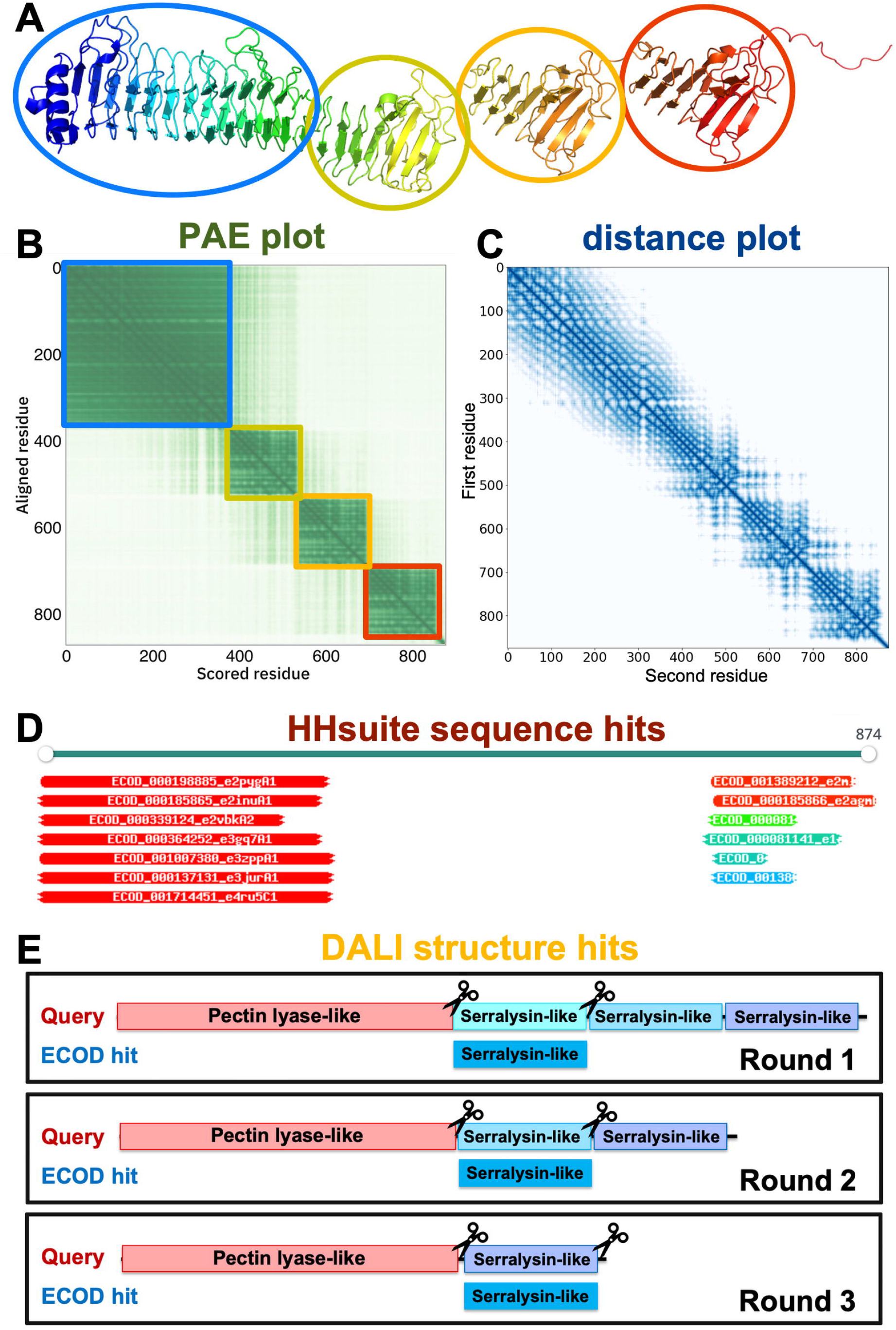
Evidence used to parse an AlphaFold model into globular domains. **(A)** An example AlphaFold model (UniProt accession: Q9ZFH0). **(B)** PAE plot for the same model. The color changes from dark green to light green as PAE value increases. Residues in the blue, yellow, orange, and red circles in **(A)** correspond to the blue, yellow, range, and red squares in **(B)** with lower PAE values inside. **(C)** Minimal inter-residue distance plot for the same model. The color changes form dark blue to light blue as distance increases. **(D)** Similar sequences detected by HHsuite. **(E)** Similar structures detected by DALI. We aligned an ECOD hit to a query in an iterative fashion to allow detection of duplicated domains in the query.

To avoid making the task easy by finding a highly similar (frequently identical) ECOD domain, we detected each protein’s closely related ECOD domains by BLAST [29]. We removed these close homologs from the HHsuite and DALI hits, respectively. We then identified “decent HHsuite hits” using two criteria: (1) the aligned residues covered 40% of the ECOD hits, (2) the HHsuite probability is at least 50%. These “decent HHsuite hits” were used for domain parsing. Similarly, we defined “decent DALI hits” as those satisfying any of the following criteria: (1) domain from the this ECOD H-group is the top hit in that region; (2) the DALI z-score between query and this hit divided by the DALI z-score when aligning this hit to itself is higher than 0.25; (3) the percentage of aligned residues in the hit domain is more than 50%; (4) the DALI z-score between query and this hit is better than the 25% quantile of DALI z-scores for comparisons of domains from the same ECOD H-group; (5) the DALI z-score between query and this hit is better than the 25% quantile of DALI z-scores for comparisons between the hit domain and other domains from the same ECOD H-group as the hit; the percentage of aligned residues in the hit domain is more than the 25% quantile of this percentage for comparisons between the hit domain and other domains from the same ECOD H-group as the hit; (6) the same hit is also detected by HHsuite. The “decent HHsuite hits” and “decent DALI hits” are not necessarily homologous to domains in the query, but we considered them to be sufficiently confident to assist domain parsing.

### Evidence to parse AlphaFold models into ECOD domains

Each AlphaFold model is accompanied by a PAE plot (**Figure 2B**) that specifies the estimated errors in inter-residue distances. PAEs reflect predicted flexibility between residue pairs. If two domains **(Figure 2A)** are connected by a flexible linker, residue pairs within the same domain are expected to show low PAEs, while pairs from different domains have high PAEs. Therefore, as AF developers also noted in AFDB, the PAE plots are suggestive of domain boundaries (**Figure 2B**). Furthermore, the PAE plots can be used to detect intrinsically disordered regions: disordered residues exhibit high PAE values except those few that are nearby in sequence. However, a PAE plot by itself is insufficient to identify evolutionary units, because two ECOD domains might be closely packed against each other, show low cross-domain PAEs, and appear as a single domain in a PAE plot. Thus, we exploited additional evidence to parse domains, including the inter-residue distances in AlphaFold models (**Figure 2C**) and similar domains in ECOD found by sequence (HHsuite, **Figure 2D**) and structure (DALI, **Figure 2E**) searches. Although these criteria were designed to parse AlphaFold models into ECOD domains, they can be generalized to other classifications if HHsuite and DALI are used against other databases.

Using our benchmark of 18,759 AlphaFold models, we used the above evidence to calculate the probabilities for a pair of residues to be located in the same domain. We binned the residue pairs by their PAEs **(Figure 3A)** and their distances **(Figure 3B)** in the 3D structure, respectively. We counted the number of residue pairs (*N*_*same*_) in the same domains and number of pairs (*N*_*diff*_) in different domains in each bin, and a residue pair in this bin will receive a “same-domain” probability of *N*_*same*_ ⁄(*N*_*same*_ + *N*_*diff*_). The probabilities derived from PAEs and distances are denoted as *P*_*PAE*_ and *P*_*DIST*_, respectively. Based on this benchmark, if two residues show PAE values of less than 8Å, the probability for them to be in the same domain is at least 50%. Similarly, if two residues are less than 35Å, the probability for them to be in the same domain is at least 50%.

**Figure 3.**
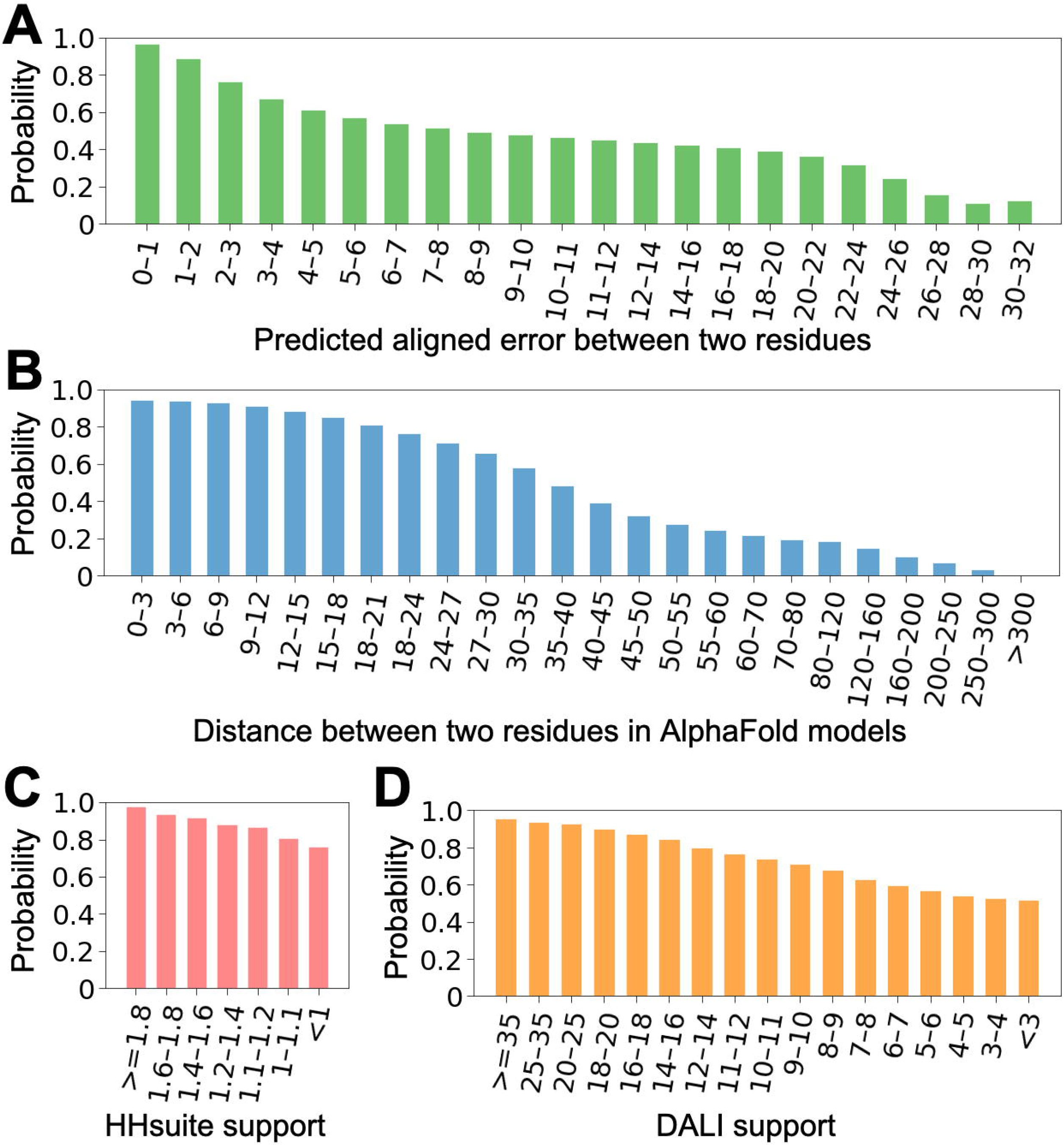
The probabilities for a residue pair to be in the same domain derived from four parameters: **(A)** PAE, **(B)** inter-residue distance, **(C)** HHsuite support, and **(D)** DALI support. The values for these parameters were binned and probability was calculated as the fraction of residue pairs to be in the same domain in each bin based on our benchmark.

Being aligned to the same ECOD domain by sequence or structural similarity provides additional support for two residues to be in the same residues, and we expect the strength of such support depends on whether the detected similarity is confident. We used the well-established confidence measurements for HHsuite and DALI, i.e., HHsuite probabilities (*HHS*_*p*_, range between 0 and 1) and DALI z-scores (*DALI*_*z*_), respectively. For a pair of residues, we identified all “decent HHsuite hits” to which both residues were aligned to, and we termed them supporting hits. We integrated the confidence of these supporting hits using the following formula to obtain the HHsuite support for a residue pair:

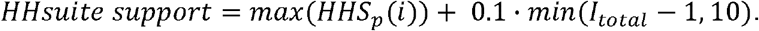

We added 0.1 · *min*(*I*_*total*_ − 1, 10) to the maximal HHsuite probability, where *I*_*total*_ is the total number of supporting HHsuite hits. Addition of this term allows a pair of residues to receive better support from HHsuite if they are simultaneously aligned to multiple HHsuite hits. Similarly, we identified all “decent DALI hits” for each pair of residues and obtained DALI support by the following formula:

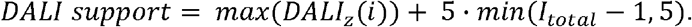

We binned the residue pairs by their HHsuite support values **(Figure 3C)** and DALI support values **(Figure 3D)**, respectively. Similar to our treatment of PAEs and inter-residue distances, a residue pair in a bin will receive the “same-domain” probability of *N*_*same*_ ⁄*(N*_*same*_ + *N*_*diff*_), where *N*_*same*_ and *N*_*diff*_ are the number of pairs from the same and different domains in this bin, respectively. The probabilities derived from HHsuite and DALI hits are denoted as *P*_*HHS*_ and *P*_*DALI*_, respectively. *P*_*HHS*_ and *P*_*DALI*_ positively corelate with HHsuite support and DALI support, respectively. In our benchmark, *P*_*HHS*_ and *P*_*DALI*_ are always larger than 0.5 based on our benchmark, because even less confident ECOD hits still primarily map to single domains. Therefore, we assigned *P*_*HHS*_ of 0.5 for residue pairs that were never aligned to the same HHsuite hit. Similarly, we assigned *P*_*DALI*_ of 0.5 for residue pairs that were never aligned to the same DALI hit.

We calculated the geometric mean of *P*_*PAE*_, *P*_*DIST*_, *P*_*HHS*_, and *P*_*DALI*_ to get the combined probabilities, *P*_*COMB*_, using the following formula:

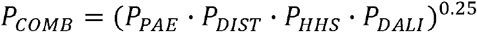

*P*_*COMB*_ was used to evaluate the probability for any pair of residues to be in the same domain, and these probabilities were used to parse domains by grouping residues with high *P*_*COMB*_ to the same domain. A visual summary of the combined principles behind DPAM is presented in Figure 4: interresidue distances, PAE measures, and consensus homologies are combined to group regions of proteins together into domains.

**Figure 4.**
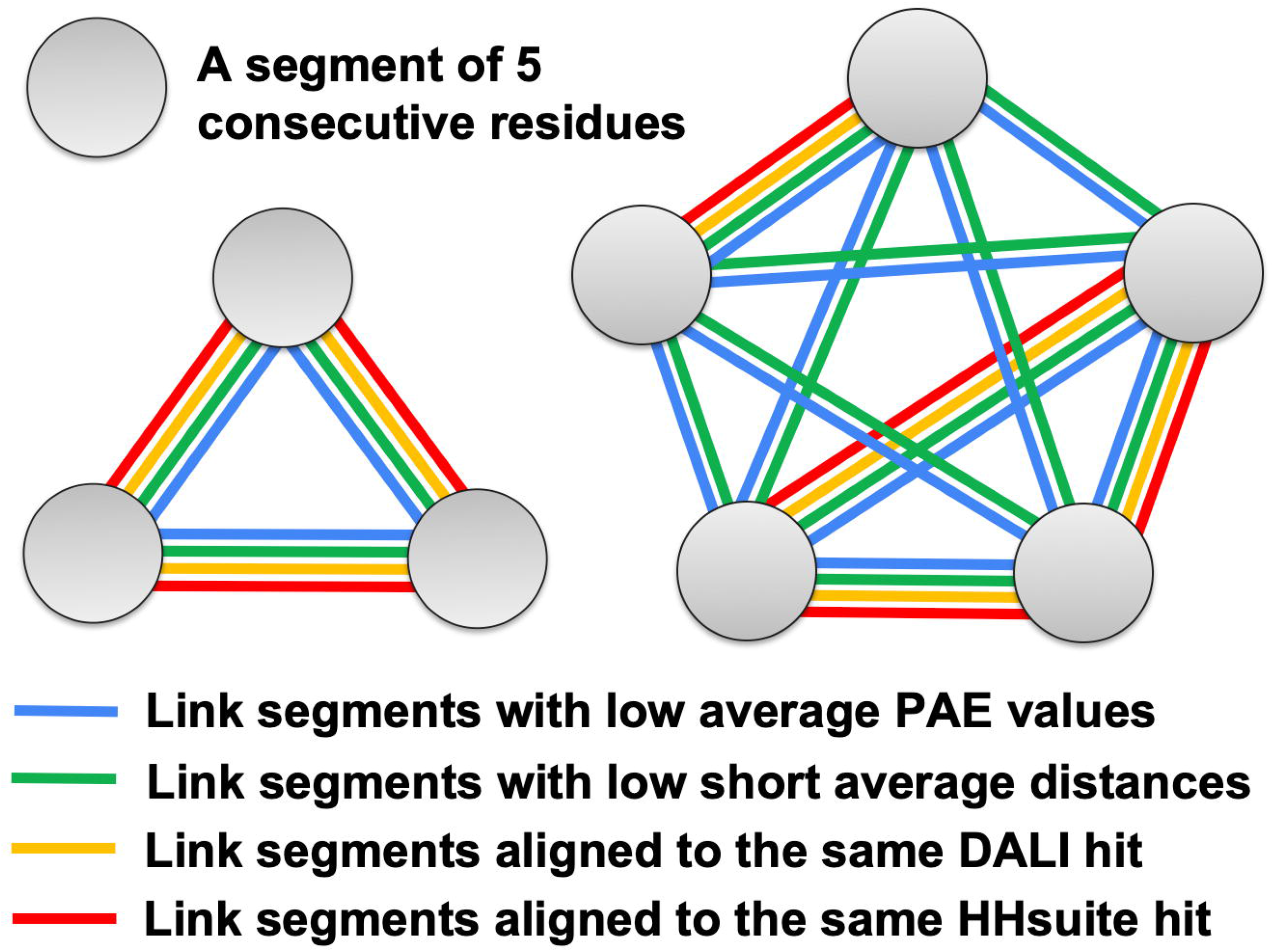
Consensus measures of prediction models govern domains identified by DPAM. DPAM relies on a consensus of PAE (blue), interreside distances (green), sequence similiarity (yellow), and structural similiarity (red) to determine domains in structural predictions.

### Clustering residues by combined probabilities

Before parsing a protein into domains, we first identified disordered regions or flexible helical domain linkers using PAE values. These regions are expected to have high PAE values against other residues in different secondary structure elements due to their flexible nature. Multiple features were used to identify such regions. First, we defined secondary structure elements in AlphaFold models aided by DSSP [30]: 3 or more consecutive residues annotated as “B” or “E” by DSSP are considered as a beta strand, and 6 or more consecutive residues annotated as “G” or “H” or “I” by DSSP are considered as an alpha helix. Second, we identified residues that were aligned to decent HHsuite or DALI hits in ECOD (excluding those from special architectures), and we dubbed them “candidate in-domain residues”: disordered segments should not contain a high fraction of “candidate in-domain residues”. Third, for each residue (target residue), we identified other residues (*C*_*PAE*<12_) that are (1) separated by at least 10 residues in sequence, (2) in a different secondary structure element from the target residue, and (3) showing relatively low PAE (<12Å) to the target residue. Residues in a disordered region or flexible helix linkers are expected to show high PAE to other residues except for nearby residues in sequence and those in the same secondary structure element (e.g, a long single helix). Using these features, we applied a sliding window of 10 residues and considered a window to belong to a disordered region or helical domain linker if its average *C*_*PAE<12*_ is no more than 3 and the number of “candidate in-domain residues” is no more than 5.

We partitioned residues in a protein to consecutive segments of 5 residues. If 3 or more residues in a segment overlapped with the disordered regions or flexible helical domain linkers, we excluded this segment. For all remaining segments, we computed the average *P*_*COMB*_ over all residue pairs, *res*_*i*_ and *res*_*j*_ for every pair of segments, *seg*_*x*_ and *seg*_*y*_ : we required *res*_*i*_ to be from *seg*_*x*_, *res*_*j*_ to be from *seg*_*y*_, and *res*_*i*_ and *res*_*j*_ to be more than 5 residues apart. We then performed clustering of the segments to group segments showing relatively high *P*_*COMB*_ into the same domain using the following procedure.

We sorted segment pairs by *P*_*COMB*_, and only considered pairs with *P*_*COMB*_ greater than *cutoff*_*P*_, a parameter to be optimized. The top-ranking segment pair was clustered into one candidate domain. Starting from the second pair, we iterated over the existing candidate domains to identify candidate domains with overlapping segments with this pair. We handled three possible scenarios. First, if neither segment in the current pair was previously included in a candidate domain, we create a new candidate domain. Second, if only one segment in this pair was present in a previously defined domain, we examined if the other segment should be merged into that domain by comparing the average *P*_*COMB*_ for segments within the domain (*intra_aP*_*COMB*_) and the average *P*_*COMB*_ between the new segment and existing segments in the domain (*inter_aP*_*COMB*_). We merge the new segment into the existing domain if *inter_aP*_*COMB*_ *· cutoff*_*M*_ *> intra_aP*_*COMB*_, where *cutoff*_*M*_ is another parameter to be optimized. Third, if the two segments in this pair were present in two previously defined domains, we examined if these two domains should be merged. We computed the average *P*_*COMB*_ for segments within the first domain as *intra*1_*aP*_*COMB*_, average for segments within the second domain as *intra*2_*aP*_*COMB*_, and average for pairs of segments coming from two different domains as *inter_aP*_*COMB*_. We merged the two existing domains if *inter_aP*_*COMB*_ · *cutoff*_*M*_ *> intra1_aP*_*COMB*_ or *inter_aP*_*COMB*_ · *cutoff*_*M*_ *> intra*2_*aP*_*COMB*_ was satisfied; *cutoff*_*M*_ is the same parameter as used in the previous scenario.

The above domain parsing routine only contains two parameters, *cutoff*_*P*_ and *cutoff*_*M*_, that need to be optimized. Since *cutoff*_*P*_ is the minimal *P*_*COMB*_ to classify two segments into the same domain, we expect its value to be slightly above 0.5. Since *cutoff*_*M*_ is the ratio between average intra-domain *P*_*COMB*_ and average inter-domain *P*_*COMB*_ when merging two candidate domains, and we expect its value to be slightly above 1. We performed a grid search to find the optimal *cutoff*_*P*_ and *cutoff*_*M*_ values: for *cutoff*_*P*_, we tested values between 0.4 and 0.7 with a step size of 0.02; for *cutoff*_*M*_, we tested values between 0.9 and 1.5, with a step size of 0.05. The best cutoffs selected using the performance evaluation routine below. The following cutoffs, 0.64 for *cutoff*_*P*_ and 1.1 for *cutoff*_*M*_, were chosen for the current version of DPAM.

After investigating all segment pairs with *P*_*COMB*_ above *cutoff*_*P*_, we obtained an initial set of domains. These initial domains were first modified to include short (≤ 20 residues) inserted segments (mostly disordered segments) that did not significantly overlap (≤ 10 residues) with other domains. Afterwards, if a domain contains multiple discontinuous segments, we excluded segments containing less than 10 residues. Finally, short domains of less than 25 residues were also removed. The current version of DPAM is available at https://github.com/CongLabCode/DPAM.

### Performance evaluation

We applied our DPAM to our assembled benchmark set. Since DPAM integrates both structure-based and homology-based evidence, we compared its performance against these two types of methods. The structure domain parsers, PDP and PUU, were applied to AlphaFold models in the benchmark set. In addition, among the “decent HHsuite hits” (closely related ECOD domains detectable by BLAST were removed), we identified a set of non-overlapping best hits among them. We first ranked these hits by HHsuite probability. Starting from the top-ranking hit, we would include a hit to this set if the majority (>75%) of the query residues it mapped to were not covered by previous hits. We call these hits “best HHsuite hits”. The query segments aligned to these “best HHsuite hits” were considered as domains based on HHsuite. Similarly, we identified a set of “best DALI hits” according to DALI z-scores, and detected domains based on DALI for each query AlphaFold model. Thus, we obtained domains by 5 methods for each model, including DPAM, PDP, PUU, HHsuite, and DALI.

We compared the parsed domains by each method against the domains defined by ECOD. Out of the 18,759 AlphaFold models in our benchmark set, ECOD currently annotated 28,348 domains. We removed 76 domains that significantly overlap (>50% residues in the domain) with the disordered regions or flexible domain linkers we detected and termed the remaining 28,272 as reference domains. For each reference domain, if over 50% of residues in it was included in the predicted domains by a method, we considered this domain to be covered by that method. The fraction of domains that were covered by different methods is shown as the blue bars in **Figure 5A**. Both PDP (98.6%) and DPAM (99.5%) domains covered most of the ECOD domains; however, the high coverage of ECOD domains by PDP is due to that PDP included a much higher fraction of residues into its domains (92.3%) than DPAM (**Figure 5B**). However, a significant fraction (6%) of residues in PDP domains belong to disordered regions or flexible linkers in AlphaFold models, and they should not be included in the globular domains.

**Figure 5.**
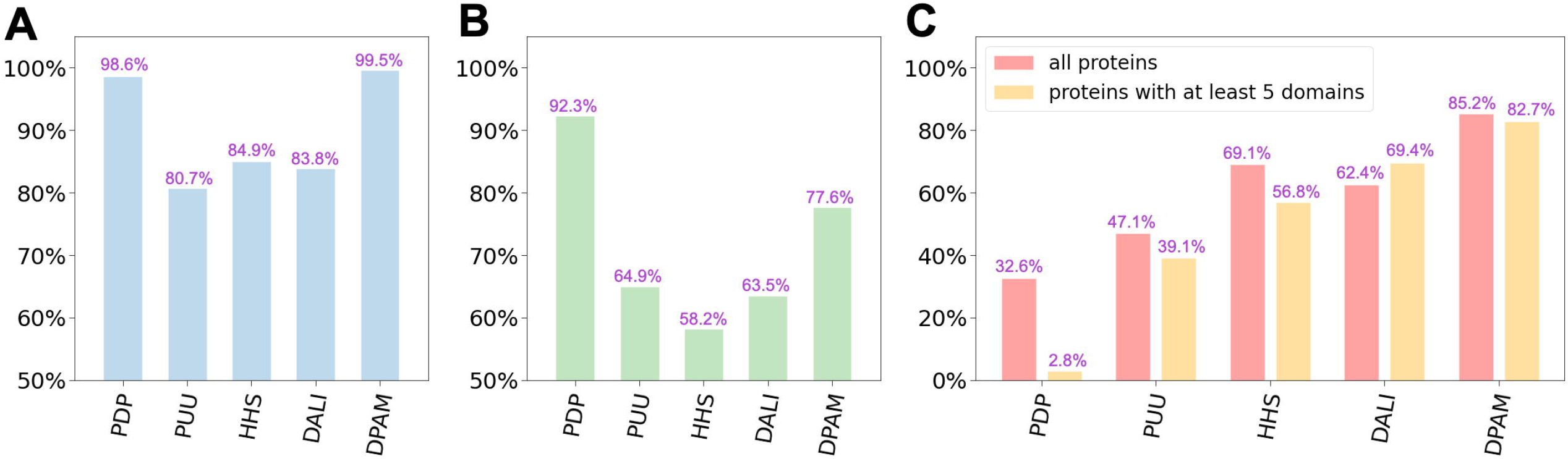
Performance evaluation of DPAM against existing structure-based domain parsers (PDP and PUU) and assignment based on similar ECOD domains found by sequence (HHS: HHsuite) and structure similarity searches (DALI). **(A)** The fraction of ECOD domains that were covered in domains annotated by different methods. **(B)** The fraction of residues that were covered in domains annotated by different methods. **(C)** The fraction of domains whose boundaries were correctly predicted by different methods.

We further determined if a reference domain’s boundary agrees with a predicted domain by a method using the following criteria: (1) the fraction of overlapping residues is higher than 75% of all residues for both the reference domain and the predicted domain; or (2) the numbers of non-overlapping residues in both the reference domain and the predicted domain are no more than 10. The fraction of domains that were correctly annotated by different methods is shown in **Figure 5C**. DPAM significantly outperformed other methods. The second-best method is HHsuite. Indeed, domain assignment by sequence similarity is efficient as long as a confident homolog could be detected, and this is how most domain assignment was derived currently in ECOD [9]. However, when a confident homolog cannot be detected by sequence, domain parsing by structural evidence is helpful. DPAM integrates both homology and structural evidence, and thus it could be particularly useful for domains cannot be easily assigned by homology.

To further evaluate the performance of DPAM, we manually studied the results for AlphaFold models containing a large number of domains (>=5). The performance of DPAM on these proteins is slightly worse (82.7%), but it is still much better than other methods (**Figure 5C**, orange bars). A number of correctly parsed models are shown in **Figure 6A-6E**. These examples suggest that DPAM can correctly parse multi-domain proteins, even in cases when domains are tightly packed against each other or proteins containing tandem repeat domains. Our manual study also revealed scenarios where DPAM tends to make mistakes. The first is when a domain is not compact and thus lack long-range contacts, such as the elongated beta-sheet in blue circles in **Figure 6F**. The second is when domains are small or poorly modeled, such as the zinc-fingers in **Figure 6G**. Zinc fingers are small domains whose folding relies on zinc ions. Due to the lack of zinc in AlphaFold models, zinc fingers tend to be poorly modeled and prone to errors in domain parsing.

**Figure 6.**
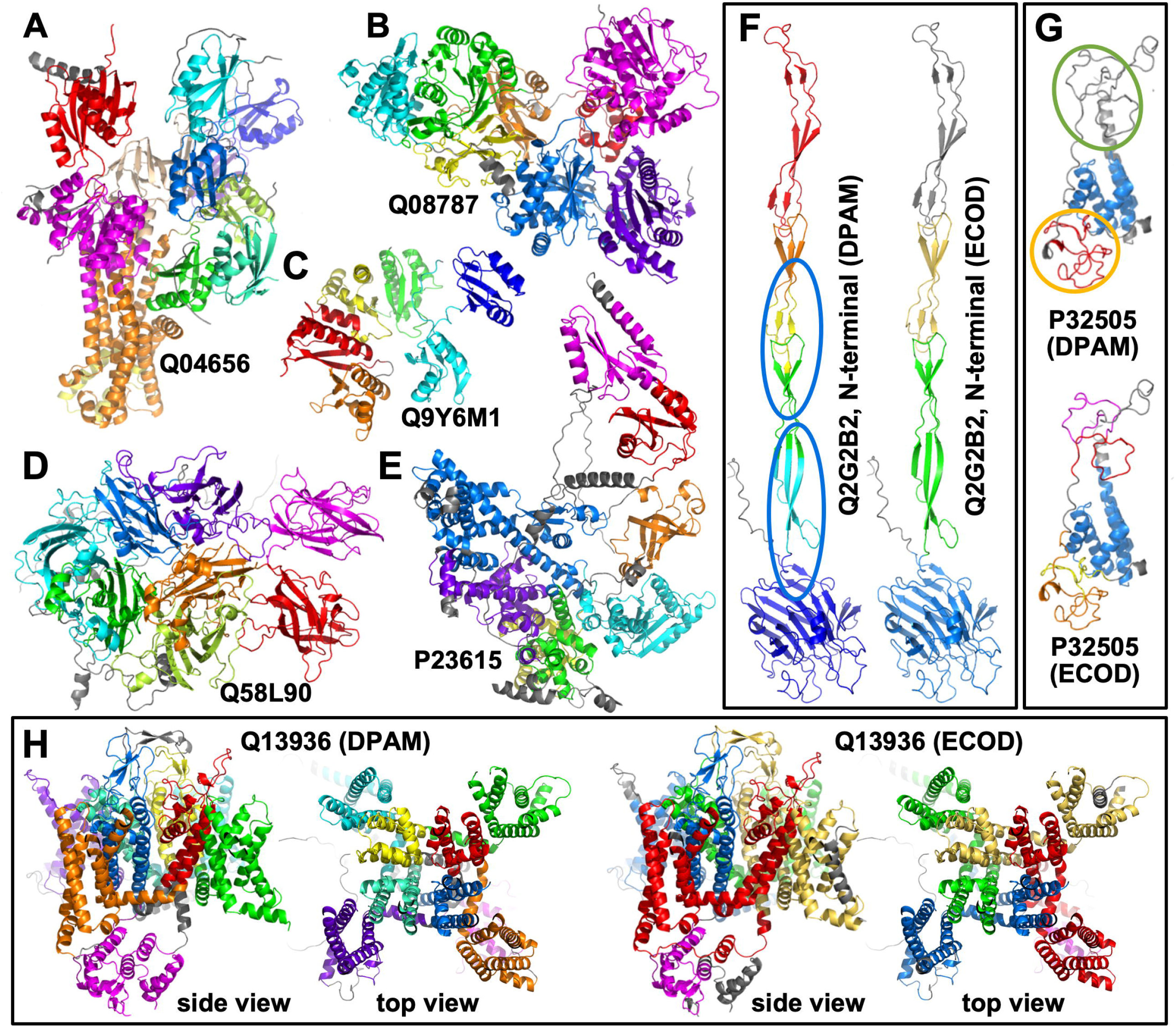
Examples of parsed domains in AlphaFold models by DPAM. Different domains in a protein are colored in differently from blue (or purple, N-terminal) through green, yellow, to red (or magenta, C-terminal). Non-domain regions are colored in grey. **(A-E)** Cases where DPAM domain definitions agree with ECOD definitions. **(F)** A case where DPAM domain definitions are not all accurate, and domains that are incorrectly split are in blue circles. (**G**) A case where DPAM missed some small zinc fingers (in a green circle) that are poorly modeled, and combined multiple consecutive zinc-fingers (in orange circles). **(H)** a case where boundaries of DPAM domains differ from ECOD domain boundaries, but the DPAM domains appear to be more meaningful.

However, in many cases, although DPAM’s domain boundaries differ remarkably from the ECOD definition, a close inspection revealed that DPAM’s domain definitions are equally or more proper. One example is shown in **Figure 6H**, the voltage-dependent L-type calcium channel subunit alpha-1C (CAC1C). CAC1C contains 24 transmembrane helices (TMHs) and adopts a 4-fold symmetry. These TMHs were parsed into 4 domains by ECOD (**Figure 6H** right), and each domain with 6 TMHs corresponds to one asymmetric unit in the 4-fold symmetric structure. Each of the 4 domains utilizes two THMs to form the central channel for calcium to go through. Because these two central TMHs are used for oligomerization between the 4-domains, they do not pack tightly against the other 4 peripheral TMHs in each domain. Therefore, DPAM considered the 2 central TMHs to form a separate domain from the 4 peripheral TMHs, a decision that is more proper from the structural perspective but less proper from the evolutionary standpoint. CAC1C contains another cytoplasmic domain (magenta in **Figure 6H**) that were both classified in ECOD and recognized by DPAM. However, DPAM assigned a more proper boundary (**Figure 6H** left) for this domain while the ECOD domain missed 2 helices (grey in **Figure 6H** right) from this domain.

## Conclusion

We developed a domain parser for AlphaFold models that combines predicted aligned errors, inter-residue distances in the 3D structures, and similar domains found by sequence and structural similarities. DPAM significantly outperforms existing structure-based domain parsers and homology-based domain assignment. Although DPAM was developed on the basis of ECOD, it can be easily extended to work with other structure classifications. We expect this tool to simplify and accelerate the classification of AlphaFold models into their evolutionary context and allow the scientific community to benefit the most from these valuable structural data.

## Acknowledgements

QC is a Southwestern Medical Foundation endowed scholar. JZ is supported by a training grant RP210041 from Cancer Prevention and Research Institute of Texas. This research is also supported by grant I-2095-20220331 to QC and I-1505 to NVG from the Welch Foundation. This research is also supported by NSF 2224128 (DBI) and NIH GM127390 to NVG.

